# Hybrid *de novo* and haplotype resolved genome assembly of Vechur cattle - elucidating genetic variation

**DOI:** 10.1101/2023.10.27.564393

**Authors:** Poorvishaa V. Muthusamy, Rajesh Vakayil Mani, Shivani Kumari, Manpreet Kaur, Balu Bhaskar, Rajeev Raghavan Pillai, Thankappan Sajeev Kumar, Thapasimuthu Vijayamma Anilkumar, N. Sadananda Singh

## Abstract

Cattle contribute to the nutritional needs and economy of a place. Performance and fitness depends on the response and adaptation to local climatic conditions. Genomic and genetic studies are important for advancing cattle breeding and availability of relevant reference genomes are essential. In the present study, the genome of a Vechur calf was sequenced on both short-read illumina and long-read nanopore platforms. Hybrid *de novo* assembly approach was deployed to obtain an average contigs length of 1.97 Mbp and N50 of 4.94 Mbp. Using a short-read genome sequence of the corresponding sire and dam, a haplotype-resolved genome was also assembled. In comparison to the taurine reference genome, we found 28982 autosomal structural variants, 16926990 SNVs with 883544 SNVs homozygous in the trio samples suggesting high prevalence of these SNPs in the population. Many of these SNPs have been reported to be associated with various QTLs including growth, milk yield and milk fat content which are crucial determinants of cattle production. Further, population genotype data analysis indicated that the present sample belongs to an Indian cattle breed forming a unique cluster of *Bos indicus*. Subsequent F_ST_ analysis revealed differentiation of Vechur cattle genome at multiple loci, especially those regions related to whole body growth and cell division especially IGF1, HMGA2, RRM2 and CD68 loci suggesting a possible role of these genes in its small stature and better disease resistance capabilities in comparison with the local crossbreeds. This provides an opportunity to select and engineer cattle breeds optimized for local conditions.

## Introduction

Cattle contribute to the nutritional needs and economy of a place. The demand for food from animal sources is rapidly increasing especially in the developing countries. According to OECD-FAO (Organisation for Economic Co-operation and Development and the Food and Agricultural Organization) of the United Nations Agricultural Outlook 2023-2032, global consumption of milk and dairy products is expected to increase by 0.8% per annum to 15.7 kg milk solid equivalents by 2032. To meet this demand, a large increase in livestock production will be required especially in developing countries. An improvement in animal feed, health and genetics will contribute mainly in achieving the target in a sustainable manner. Adopting new technologies or customization of existing technologies is being carried out in many countries. In general, cross-breeding between a highly adapted but with low productivity indigenous breed and a poorly adapted but highly productive exotic breed and further selection is conducted to develop a high yielding well-adapted crossbreed. Cross-bred under better management has shown manifold increase in milk yield thereby leading to substantial increase in household income and reduction of greenhouse gas emission. In recent times, application of genomics is becoming increasingly helpful and important for implementation of meticulously planned breeding programmes for breed improvement exercise including breed composition assignment. Genomics based approaches have been successful in developing economical genotyping panels and/or assays for use during genomic selection including ancestry proportion determination which is important during breed selection [1, 2]. Genomic information is used in genomic selection which helps in more accurate prediction of phenotypes at young age, utilization of information available for distant breeds, reducing cost, time and number of crosses as compared to traditional breeding methods [3]. Recently, climate change has led to increased incidences of higher intensity heat waves which leads to another challenge to the cattle breeding efforts as adaptation to heat stress leads to lower efficiency of production and, thus, is unfavorable to the goal of reducing GHG [2]. Indicine breeds are known for their resistance to drought, better tolerance to heat and sunlight [4] and disease resistance [5] Thus, crossbreeding with indicine breeds and genomic selection approach offers high potential to achieve yield improvement goals. However, the lack of genome sequence of indicine cattle becomes a limiting factor in carrying out genomic based breeding using indicine breeds.The only available previous reference-based genome assemblies of B. indicus cattle (Nellore breed) and other indicine breeds were using short-read sequencing platform. For B. indicus cattle (Nellore breed), reference based assembly was performed using SOLiD sequencing platform with very short read lengths of 25 and 50 bases. The recent reference-based assembly was carried out using illumina platform with readlengths of 150 bases [6,7] while there is a high quality reference genome for taurine cattle breeds [8]. Therefore, a quality reference genome of the indicine breed is still missing.

The biological and economical output efficiency is very important for the dairy farmers and it has been reported that lighter cows provide a comparatively higher economic value on land basis [9]. It has also been reported that feed efficiency (milk yield per kg feed) was negatively correlated ranging from -0.18 for wither height to -0.33 for body weight with body weight and the body measurements ranging from -0.18 for wither height to -0.33 for body weight [10]. Thus, an indicine breed with known dairy history and small size would be an important one to study or use for crossbreed development by genomic selection. There are around 75 breeds of indicine cattle majorly split between African breeds and Indian breeds. According to the animal genetic resource portal (https://nbagr.icar.gov.in/en/registered-cattle/), there are 53 registered cattle breeds in India. There are phenotypic variations among these breeds. Vechur breed found in the south-western state of Kerala where cross-breeding with taurine breeds of cattle is practiced over the last six decades to improve milk production, is a small sized, well adapted cattle with an average weight of about 133.6 ± 3.7 and 173.5 ± 6.8 kg and a height of 89.0 ± 0.7 and 99.8 ± 1.4 cm for cows and bulls, respectively. This breed was the most popular dairy breed with 2-3 liter/day in the region before it was replaced by high milk yielding crossbreeds [11]. These cattles are also well-known for their resistance to viral, bacterial, and parasitic diseases compared to the exotic cattle and their crossbreds [12,13].

In the present study, we have collected a family trio (sire, dam, and calf) of vechur cattle and the calf genome was sequenced using both short-read illumina and long-read nanopore platforms to assemble a genome using hybrid de novo assembly approach. Using short read sequences of the sire and dam, a haplotype resolved genome was also assembled. Further, genetic variants were analyzed using the taurine breed reference genome to find association with various QTLS. F statistics analysis was carried out using the new genome sequence data and other available genotyping data to find genetic loci that may differentiate vechur from rest of the indicine breed and may explain its short stature too.

## RESULTS

### Samples and sequencing

Blood DNA samples of a family trio consisting of dam (MT435), a sire (MT436), and its calf (MT434) were collected and sequenced on a short read illumina sequencing platform. The calf DNA was also sequenced using nanopore long read sequencing platform. The sequencing details are given in table 1.

**Table 1:** Details of sequencing.

| Short Read Illumina platform sequencing details |  |  |  |  |  |  |
| --- | --- | --- | --- | --- | --- | --- |
| Sample Name | Average Read length (bp)X2 | #Raw Reads (Forward /Reverse) | #Total Reads | Raw data (bp) | %GC | Coverage |
| MT434 (calf) | 151 | 449742276 x2 | 899,484,552 | 135,822,167,352 | 44 | 50.3 |
| MT435 (dam) |  | 432569400 x2 | 865,138,800 | 130,635,958,800 | 44 | 48.4 |
| MT436 (sire) |  | 559344678 x2 | 1118689356 | 168,922,092,756 | 44 | 62.6 |
| Long Read Nanopore platform sequencing details |  |  |  |  |  |  |
| Sample Name | Mean Read Length (bp) | # Total Reads (Single end) | Raw data (bp) | %GC | Coverage |  |
| MT434 | 4273 | 37,211,638 | 159,011,500,107 | 43.73 | 58.89 |  |

### De novo Genome assembly

De-novo hybrid assembly was performed for the sample MT434 (calf) using CLC Genomics workbench 22.0.5 using both nanopore and illumina reads. For this sample, sequencing from both Illumina and Oxford Nanopore NGS platforms generated raw data of 135 Gb and 159 Gb respectively, corresponding to 50.3x and 58.89x coverage respectively. It was performed in two steps as depicted in figure 1A: (i) de novo assemble a genome using long, error-prone reads, (ii) improve the de novo assembly from long reads by polishing with short, high-quality illumina reads. The refined assembly results in a genome of 2,693,805,279 bp and the assembly statistics are given in Table 2. To assess the genome completeness further, the Benchmarking Universal Single-Copy Orthologs (BUSCO) [14] was used, which has a predefined and expected set of single-copy marker genes as a proxy for genome-wide completeness. The assembled genome was used and the genome mode was selected, for lineage, Eukaryote was selected to run just on eukaryote trees to find optimum lineage. The results have been summarized in table 2.

**Figure 1:**
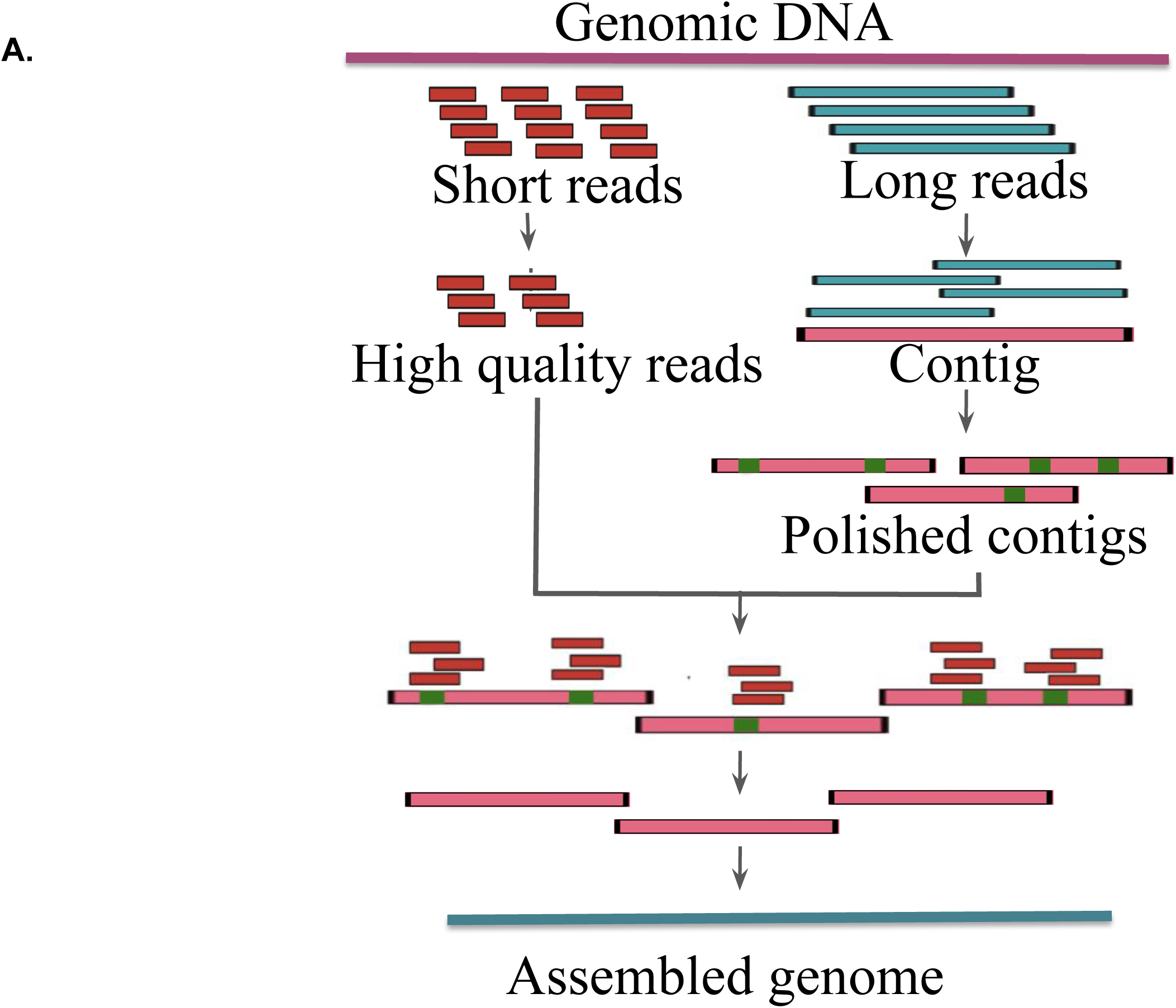
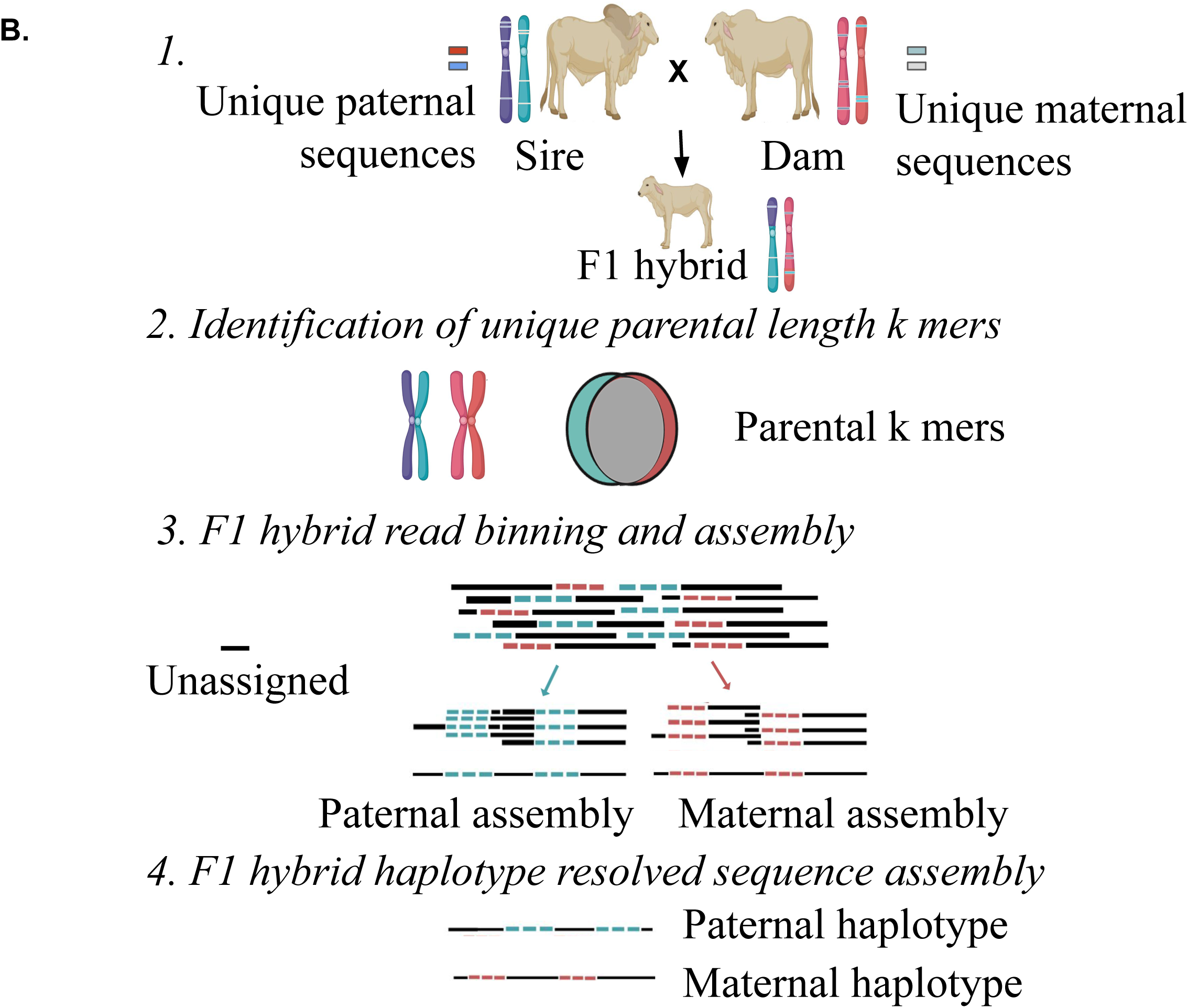
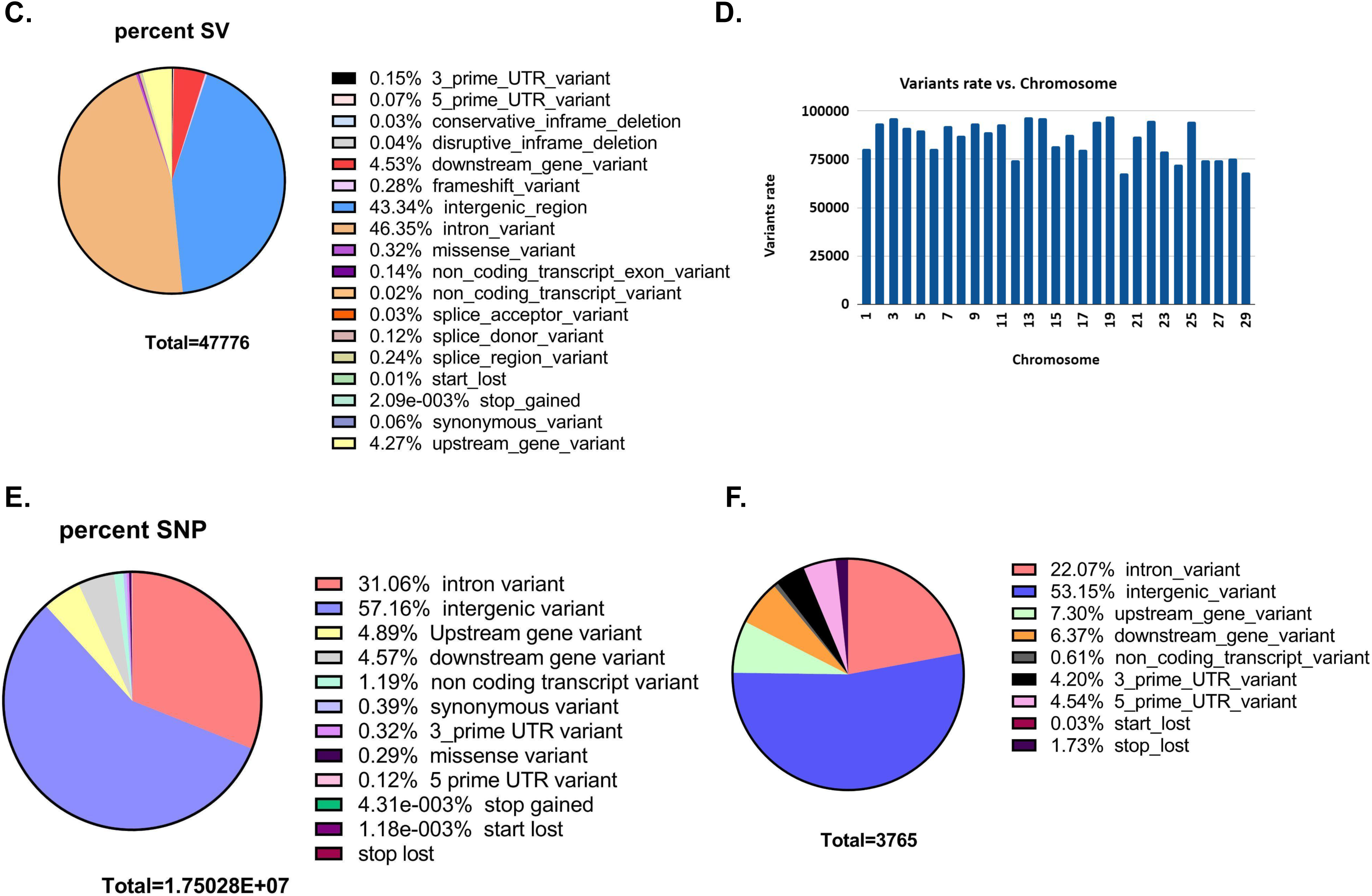
**A.**Schematic diagram showing the hybrid de novo genome assembly pipeline **B.**Schematic diagram of the trio-binning haplotype resolved genome assembly **C.** Pie chart showing per cent of structural variants with each predicted consequences of sample MT434 (calf) obtained using the ensembl variant effect predictor (VEP) **D.** Pie chart showing per cent of single nucleotide variants (SNVs) with each predicted consequences of sample MT434 (calf) obtained using the ensembl variant effect predictor (VEP) tool. **E.** and **F.** Histograms showing the number of SNVs and rate of SNVs detected on each chromosome respectively

**Table 2:** Assembly statistics and BUSCO analysis summary.

|  | **Sample:MT434** |
| --- | --- |
| **Contigs** | 1,367 |
| **Minimum Length** | 10,281 |
| **Maximum Length** | 27,791,687 |
| **Average Length** | 1,970,596 |
| **N50** | 4,946,819 |
| **N90** | 1,163,082 |
| **Total** | 2,693,805,279 |
| **BUSCO analysis** |  |
| Busco Summary | C:90.4%[S:85.1%,D:5.3%],F:2.3%,M:7.3%,n:303 |
| Complete BUSCOs (C) | 274 |
| Complete and single-copy BUSCOs (S) | 258 |
| Complete and duplicated BUSCOs (D) | 16 |
| Fragmented BUSCOs (F) | 7 |
| Missing BUSCOs (M) | 22 |
| Total BUSCO groups searched | 303 |

### Haploid resolved assembly

Haplotype resolved assemblies were generated using the trioCanu module of the Canu assembler [15]. To enable haplotype-resolved assembly of the calf, we performed short read sequencing of the dam and sire using illumina platform with a coverage of 32.58x and 37.67x respectively. These reads were quality trimmed and filtered. Haplotype binning (trio binning) was done which takes the short reads from the parental genomes to partition long reads from the offspring into haplotype-specific sets as depicted in Figure 1B. Details of the binned reads are summarized in table 3. Using the binned reads, each haplotype was then assembled independently using Long Read Support (beta) plugin of CLC Genomics workbench 22.0.5. These resulted in a paternal haplotype assembly of 2,556,074,938 bp with N50 of 1.4 Mbp and a maternal haplotype assembly of 2,618,152,939 bp with N50 of 2.0 Mbp as summarized in Table 4.

**Table 3.**
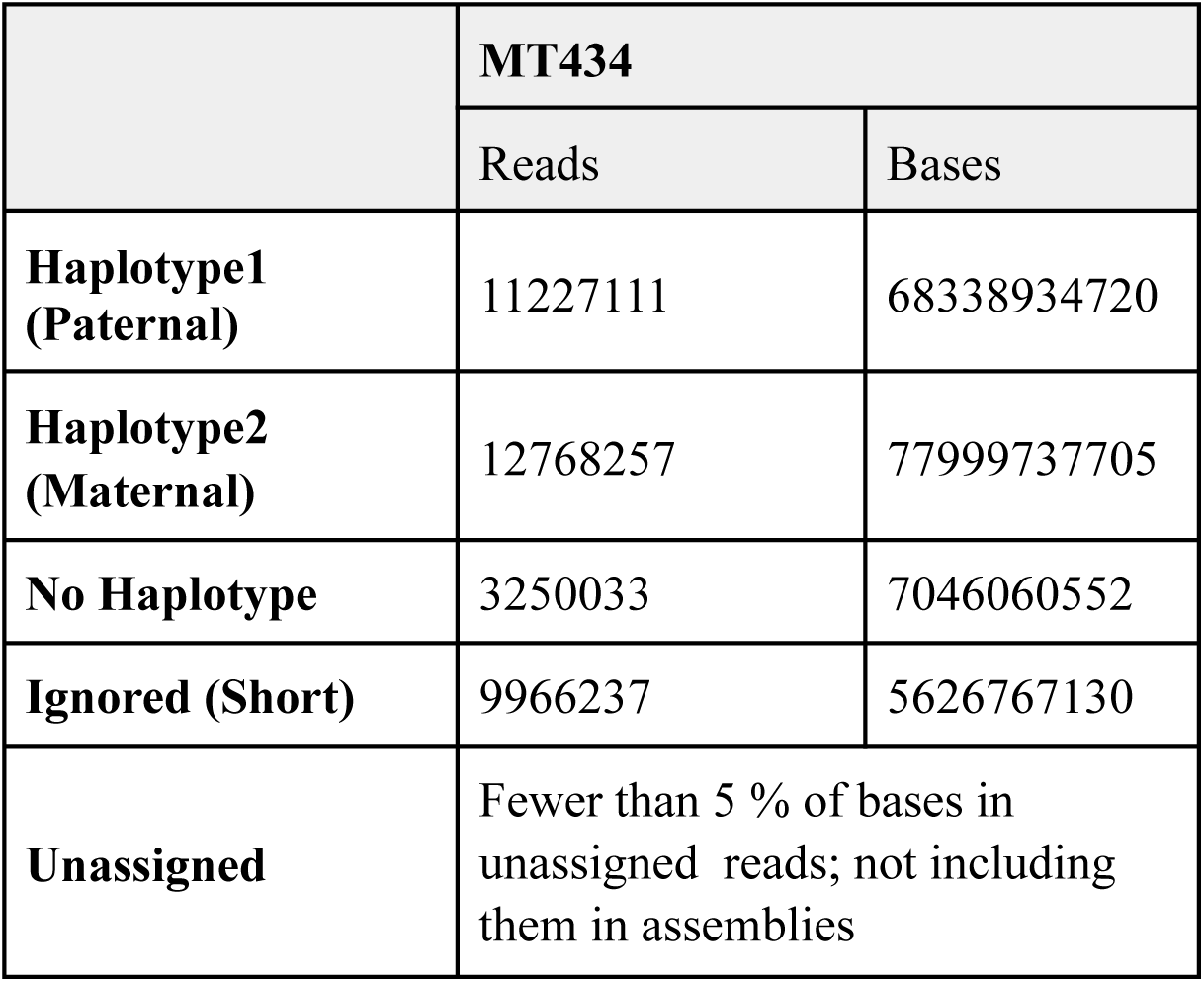
Summary of haplotype binning (trio binning)

**Table 4.** Summary of haplotype resolved assemblies.

|  | **MT434_Haplotype1** | **MT434_Haplotype2** |
| --- | --- | --- |
| **Contigs** | 4,217 | 2,956 |
| **Minimum Length** | 10,055 | 10,035 |
| **Maximum Length** | 10,270,387 | 10,894,072 |
| **Average Length** | 606,136 | 885,708 |
| **N50** | 1,373,887 | 2,074,774 |
| **N90** | 300,353 | 438,793 |
| **Total** | 2,556,074,938 | 2,618,152,939 |

### Structural and single nucleotide variant analysis

In comparison to the taurine reference genome ARS-UCD1.2.15, we detected 30434 structural variants with 28982 autosomal structural variants ranging from 50 bp to 9.97 kbp, with an average one structural variant for every 86,363 bp with mutation rate on chromosome 20 (Figure 1D). Most of the variants (∼90%) are in the intergenic and intronic regions while 529 variants (∼1.2%) are in the coding regions (Figure 1C). In addition, there are 16926990 single nucleotide variants with 1521747 novel and 15405243 existing variants. 5634648 (33.3%) in coding region, 51390 missense, 754 nonsense variants and 11 readthrough variants (Figure 1E). When analyzed using CNVnator 4.0 [16], we also detected 3395 copy number variations (2470 deletions and 925 duplications), and VEP analysis predicts 17% feature truncation, 4% feature elongation, and 8% in the coding region, as seen in Figure 1F. The PANTHER database was used to functionally annotate the 712 genes found in the inferred CNV regions, and the most enriched pathways were the IGF pathway-mitogen activated protein kinase kinase/MAP kinase cascade, T cell activation, Gonadotropin releasing hormone receptor pathway, and Interleukin signaling pathway.

We also called the variants of the dam and sire in comparison to the taurine reference genome to enquire if there are variants which may give direction in understanding different quantitative traits. We have analyzed the variants and sorted those variants which are present in homozygous condition in both the dam and sire. The homozygosity of these variants suggest the high prevalence in the vechur population. 67301 SNPs were present in homozygous condition and were further analyzed for their association with quantitative traits using the cattle QTL database (release 50) available at https://www.animalgenome.org/cgi-bin/QTLdb/BT/index. It is worthwhile to mention that many of the milk related traits and body weight were among the top quantitative traits associated with maximum number of SNVs as listed in Table 5.

**Table 5:** List of top 10 QTLs associated with homozygous SNVs.

| Trait | No. of QTL locus |
| --- | --- |
| 1. Scrotal circumference | 8468 |
| 2. Age at puberty | 8460 |
| 3. Milk fat percentage | 3546 |
| 4. Milk kappa-casein percentage | 3103 |
| 5. Percentage normal sperm | 2968 |
| 6. Milk fat yield | 2560 |
| 7. Body weight | 2246 |
| 8. Milk protein percentage | 2012 |
| 9. Milk glycosylated kappa-casein percentage | 1793 |
| 10. Calving ease | 1731 |

### Population analysis

Vechur cattle is well known for its disease resistance, better adaptation for tropical extreme climates and small stature. Classical multidimensional scaling (MDS) based on pairwise identical-by-state (IBS) distance was performed to understand or validate genetic relatedness and population stratification (i) between Vechur and different breeds of cattle in the world using the data published earlier [17], (ii) between vechur and other indian indicine breeds using the data published earlier [18]. As depicted in figure 2A, Vechur (Ind_VC) and the present trio (blue squares) cluster together with the other indian indicine breed whereas African zebu (AZs in figure 2A) breeds and taurine breeds (ETs in figure 2A) form the other two clusters. Further resolution of the MDS analysis among the indian indicine breeds, vechur (ind_VCs, purple circle in figure 2B) alongwith the trio (MTxxx, blue squares in figure 2B) forms a unique subcluster as depicted in figure 2B indicating the presence of a unique selection genetic feature. Admixture analysis also supports the above observation as shown in Figure 2C.

**Figure 2:**
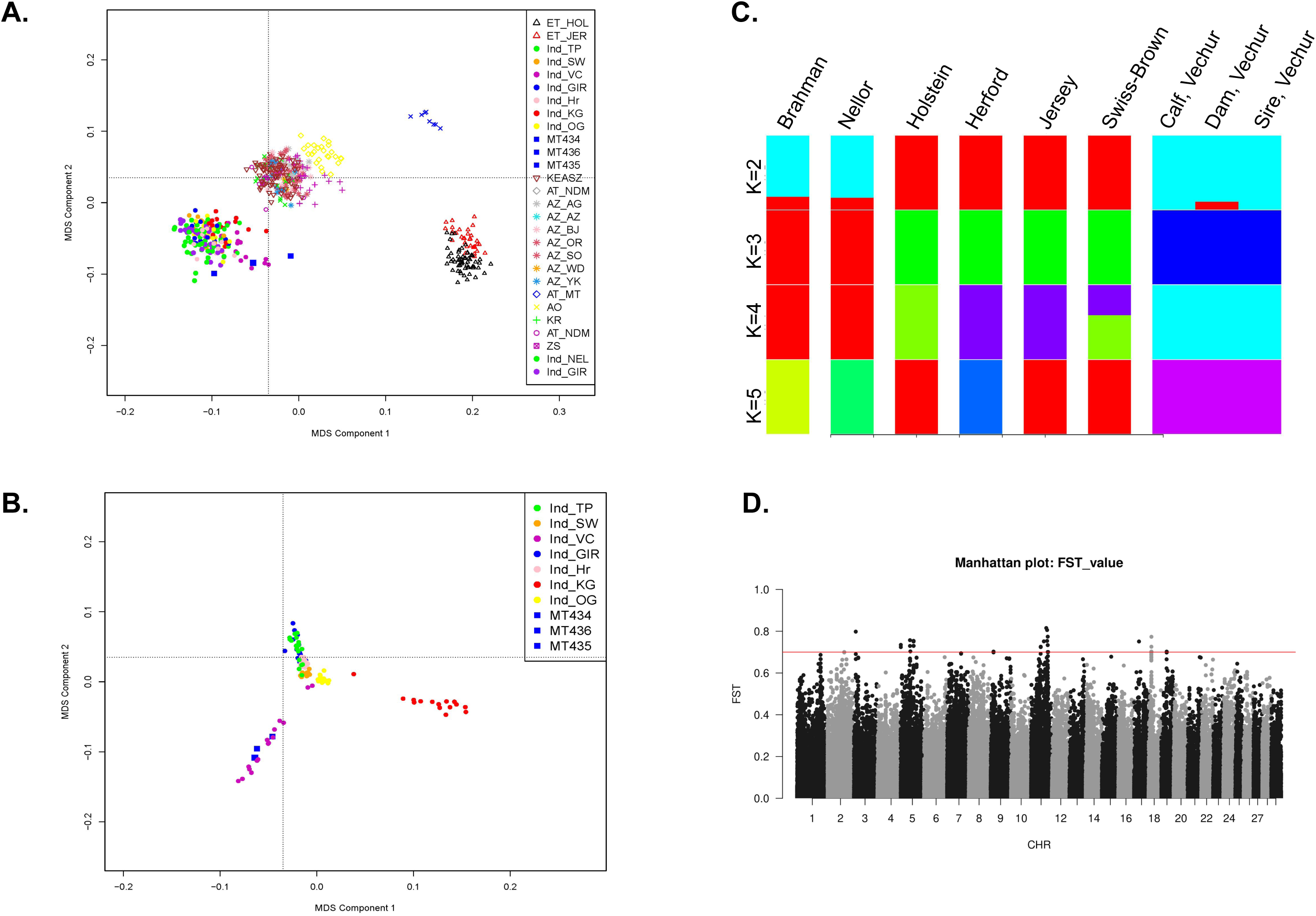
**A.** Multi-dimensional scaling (MDS) analysis of various cattle breeds including indicine breeds to show the clustering of Vechur cattle with the indicine breeds Breed abbreviation as follows: (i) ET_XXX: European taurine breeds, (ii) Ind_xxx : Indian Indicine breeds, (iii) MT434/435/MT436 are the trio vechur being used in this study, (iv) KEASZ: Kenya Small EASZ, (v) AT : African taurine, (vi) AZ_xx : African zebu breeds, (vii) AO : Sanga, (viii) KR : Uganda large EASZ and (ix) ZS : Uganda small EASZ. **B.** Multi-dimensional scaling (MDS) analysis of various indian cattle breeds showing the formation of separate cluster of Vechur cattles Breed abbreviation as follows : Ind_TP : Tharparkar, Ind_SW : Sahiwal, Ind_GIR : Gir, Ind_OG : Ongole, Ind_VC : Vechur, Ind_Hr : Hariana and Ind_KG : Kangayam.. **C.** Admixture analysis of six cattle breeds ranging from K=2 to K=5. Breed abbreviation as follows: BRM01, Brahman; NEL01, (Nelore) Indicus; HOL01, Holstein; TAU01, Hereford; JER01, Jersey; MT434 (Calf); MT435 (Dam); and MT436 (Sire) of Vechur breed. **D.** Manhattan plot of genome-wide FSTvalues (cut-off value>0.7) comparing vechur cattle vs rest of the indian breeds.

Fixation index (F**_ST_**) tests were performed to identify SNPs which are highly differentiated in vechur as compared to other indian cattle breeds. This analysis would likely reveal SNPs or genomic regions which are involved in controlling body size. As reported earlier, F_ST_ analysis results show 35 SNPs (listed in Table 6) with high F_ST_ values (>0.7) clustered mainly in certain regions of chromosome 5, 11 and 18, (Figure 2D) and house protein coding genes: IGF1, HMGA2, SRGAP1, APOB, ENSBTAG00000020828,RRM2, ZNF276 and CD68 (listed in Table 7). IGF1 is involved in growth and dysfunction of HMGA2 results in autosomal dominant growth retardation phenotype [19]. HMGA2 regulates IGF2 [20]. Using the sequence data of the present family trio that has been generated during this study, variants were called for these chromosomal locations listed in table 7. 2324 variants were detected with 520 being homozygous in all the members of the family with 27 missense (22 in APOB and 1 each in HMGA2, RRM2, IGF1 and SRGAP1 and ENSBTAG00000048587), 5 splice site variants (2 in APOB, 1 each in RRM2, SRGAP1 and HMGA2), 4 5’UTR and 13 3’UTR (1 in IGF1, 6 in RRM2 and 6 in ENSBTAG00000048587).

**Table 6:** List of SNPs (in the HDbeadchip) with F_ST_ values >0.7.

| CHR | POS | WEIR_AND_COC<br>KERHAM_$F_{ST}$ | Gene | CHR | POS | WEIR_AND_COC<br>KERHAM_$F_{ST}$ | Gene |
| --- | --- | --- | --- | --- | --- | --- | --- |
| 2 | 86180784 | 0.700374 |  | 11 | 82332335 | 0.753475 |  |
| 3 | 9132647 | 0.798089 |  | 11 | 65570076 | 0.751379 |  |
| 5 | 48008400 | 0.75717 |  | 11 | 85454517 | 0.733433 |  |
| 5 | 66488531 | 0.753664 |  | 11 | 49393563 | 0.725848 | ENSBTAG00<br>000020828 |
| 5 | 66499710 | 0.753664 |  | 11 | 49361395 | 0.723943 |  |
| 5 | 66545432 | 0.750791 | IGF1 | 11 | 87466344 | 0.700458 |  |
| 5 | 576396 | 0.734675 |  | 11 | 87472407 | 0.700458 | RRM2 |
| 5 | 66547981 | 0.729853 | IGF1 | 11 | 85772933 | 0.700374 |  |
| 5 | 66552462 | 0.729853 | IGF1 | 11 | 85782922 | 0.700374 |  |
| 5 | 66557413 | 0.729853 | IGF1 | 11 | 85789132 | 0.700374 |  |
| 5 | 66562687 | 0.729853 | IGF1 | 17 | 25405683 | 0.750791 |  |
| 5 | 48057883 | 0.729551 | HMGA2 | 18 | 14760377 | 0.773436 |  |
| 5 | 540754 | 0.723949 |  | 18 | 14647288 | 0.725848 | ZNF276 |
| 5 | 49954556 | 0.703359 | SRGAP1 | 18 | 14688492 | 0.702313 |  |
| 9 | 14680303 | 0.702501 |  | 18 | 14692912 | 0.702313 |  |
| 11 | 77988513 | 0.815351 | APOB | 18 | 14327200 | 0.701693 |  |
| 11 | 82338187 | 0.80554 |  | 19 | 27921934 | 0.702313 | CD68 |
| 11 | 85600438 | 0.774 |  |  |  |  |  |

**Table 7:** Summary of the list of all loci differentiated in the Vechur breed.

| Chromosome | Co-ordinates | Gene |
| --- | --- | --- |
| 5 | 5:66532877-66604734 | Insulin like growth factor 1 (IGF1) |
| 5 | 5:48053846-48199963 | High mobility group AT hook2 (HMGA2) |
| 5 | 5:49812166-49969012 | SLIT ROBO Rho GTPase activating protein 1 (SRGAP1) |
| 11 | 11:77953380-78040118 | Apolipoprotein B (APOB) |
| 11 | 11:49390250-49396081 | ENSBTAG00000020828 |
| 11 | 11:87466581-87473817 | Ribonucleotide reductase regulatory subunit M2 (RRM2) |
| 18 | 18:14635302-14649693 | Zinc finger protein 276 (ZNF276) |
| 19 | 19:27921927-27923997 | CD68 |

## DISCUSSION

A haplotype-resolved genome of an indicine breed has been assembled in this study. There is a significant improvement of the indicine cattle genome as compared to the presently available reference genome as reported earlier in [6] and recently built short read sequencing based genome [7]. *The use* of relevant reference genomes is important and could have a large impact on studies, especially on detecting signatures of selection as has been reported earlier [21].

In genetic analysis, use of a single taurine reference genome may generate systemic errors as was observed in case of Homo sapiens [22]. Among 53 cattle breeds of India listed at https://nbagr.icar.gov.in/en/registered-cattle/ Vechur is one of the smallest indicine breeds in the world with exceptional adaptation to the tropical weather conditions. Thus, this genome would help in unraveling genetic factors involved in such adaptation.

MDS plot and admixture analysis revealed that vechur is one of the indicine breeds and its haplotype resolved genome would serve as a better reference genome for the local and pan indian indicine breed. Most of the dairy cattle breeds in India are crossbreeds between taurine breeds like jersey and indian breeds. Availability of an indicine breed reference genome would help in genetic studies related to milk production and local environment adaptation phenotypes using state of the art genomic selection procedures. Moreover, better understanding of genetic factors may help in applying targeted genome editing technologies to introduce desirable trait related genetic variants in the genome.

We also found high genetic differentiation in multiple regions of vechur genome as compared to the other indicine breed. These regions host genes including IGF1, HMGA2, SRGAP1, APOB, ENSBTAG00000020828,RRM2, ZNF276 and CD68. IGF1 is a known growth promoting gene and reported to contribute 30-45% of growth in mice [23, 24]. HMGA2 deficient mice, zebrafish and horse also show reduced growth [25, 26]. The HMGA2 deficient phenotype of reduced growth may be explained by its regulation of IGF2 [20] which is again related to IGF1. There are 1 missense IGF1 variants (T151M) and HMGA2 (G41C) variants which are homologous in all the members of the family suggesting a high prevalence of these variants in vechur cattle. These and other variants in these genes are likely to contribute majorly in the small stature phenotype of this cattle breed. CD68 is a macrophage marker and reported to be involved in inflammatory reactions [27]. We believe this Vechur genome assembly will provide genomic resources for evolutionary studies in combination with the other bovine species. Overall, a haplotype-resolved genome of an indian indicine cattle is reported in this study and will help in genomic selection studies related to improved milk yield, improved efficiency, and better adaptation.

## MATERIALS AND METHODS

### DNA isolation from blood samples

2 ml of blood samples were taken in a 15 ml falcon tube and 4 ml of chilled lysis buffer (150 mM NH_4_Cl, 10 mM 1M KHCO_3_ and 0.1 mM EDTA) was added. It was kept on ice for 10 min after mixing. It was then centrifuged at 7000 rpm for 10 min at 4 °C. The supernatant was discarded and the process was repeated until pellet is clear of RBC (2-3 washes are sufficient). 300 µl of extraction buffer (400mM NaCl, 2 mM EDTA, 10 mM TrisCl pH8.0) was added and mixed well. 100µl of Proteinase K (0.2mg\ml) and 125µl of 20% SDS was added, mixed and incubated at 56 °C for six hours or overnight. Phenol chloroform extraction was performed by adding 500µl of Phenol-Chloroform-Isoamylalcohol (25:24:1) to the mixture and mixed well by gently inverting the tube up and down for 10 min to get a milky emulsion. Centrifuged at 10000 rpm for 6mins and the upper aqueous phase gently was extracted again with 500µl of Chloroform-Isoamylalcohol (24:1). The DNA was precipitated by adding 1/10th volume of 3M sodium acetate (of aqueous layer) and 2.5times volume of chilled absolute alcohol followed by centrifugation first at 10,000 rpm for 5 min, then at 12,000rpm for next 5 min and last 14000 rpm for 10 min at 4°C. The pelleted DNA was washed 2 times with 300µl of 70% ice cold ethanol and dried at room temperature. It was then dissolved in 100µl nuclease free water or 1XTE buffer by incubating at 56°C for 10min. The DNA was then stored at -20°C until further use.

### Sequencing

Extracted DNA was sequenced on both the Illumina and Oxford Nanopore platforms. The short reads produced by Illumina technology were used to estimate genome size and correct errors in the assembled genome. Long readings from the Oxford Nanopore device, on the other hand, were used in the actual genome assembly process. A library with insert lengths of 150 base pairs was sequenced on the Illumina Nextseq technology to achieve this goal. In addition, another library with an average length of 20 kilobases was created using the Oxford Nanopore platform in line with the manufacturer’s instructions.

### Genome assembly

The “De Novo Assemble Long Reads” tool within CLC Genomics Workbench version 22.0.5 was used with a specialized plugin for de novo hybrid assembly. This tool is designed for processing long, error-prone reads, like those from Oxford Nanopore Technologies. It utilizes open-source components: minimap2, miniasm, raven, and racon. The hybrid assembly involves two main steps: first, the de novo assembly of a genome using long, error-prone reads,, and second, the refining of the initial de novo assembly produced from long reads using short, high-fidelity reads.

The Uncorrected nanopore reads were used directly. The process begins with finding overlap alignments among the input reads using miniasm/minimap2. These overlaps are pre-processed with pile-o-grams, creating an assembly graph, which is then simplified to produce contigs using raven assembler. The Default settings (k = 15, w = 5, minimum contig size = 1000) and two rounds of racon polishing were applied. Contig polishing is performed twice using racon/minimap2, which improves a partial order alignment (POA=500) of the reads against the contigs and contig quality through rapid consensus calling.

The assembly was further polished with high-quality Illumina short reads using racon and enhancements from minipolish . Racon uses a divide-and-conquer strategy for rapid consensus calling. Trimmomatic 0.39 was used to trim and filter Illumina reads for quality and length. These reads were then mapped to assembled contigs to refine them. Most contigs had roughly 40x coverage or higher. The binned reads for individual contigs were retrieved and used for polishing. The partial order alignment (POA) window was set to 500 bp, and the minimum sequence length for output was 10,000 bp, as all contigs were longer. The remaining settings remained consistent.

### Haplotype resolved assembly

Haplotype resolved assemblies were also made using the trioCanu module of the Canu assembler [15]. Prior to assembly, haplotype binning (trio binning) was done which takes the short reads from the parental genomes to partition long reads from the offspring into haplotype-specific sets. Each haplotype is then assembled independently, resulting in a complete diploid reconstruction. For MT431, the parental reads MT432-dam & MT433-sire were quality trimmed & filtered and then are used for trio binning using the long reads of their offspring MT431. Similar approach was used for MT434, where the QC passed parental short reads MT435-dam & MT436-sire are used for binning the long reads of MT434 (offspring). The trio binning divides the total reads into paternal and maternal groups on the basis of the presence of the haplotype-specific k-mers in those bins. These haplotypes were then assembled using the Long Read Support (beta) plugin of CLC Genomics workbench 22.0.5.

### Structural Variant analysis

The initial draft assembly was aligned using nucmer (l=100, c=500) against the reference genome Bos taurus (cattle) - Hereford breed (ARS-UCD 1.2; GCF_002263795.1) to obtain a delta file, which was then uploaded to Assemblytics to analyze alignments. The input file (OUT.delta.gz) has been provided for loading on the Assemblytics web server and can be used to view the results dynamically. A Ensembl Variant effect predictor [25] was used to predict the consequences of the structural variants.

### Alignments and variant identification

Prior mapping, adapter sequences and low-quality reads were removed using Trimmomatic 0.39 and high-quality reads were aligned to the UMD3.1 bovine reference genome assembly using the bwa-mem option of Burrows-Wheeler Alignment programme (BWA) version 0.7.5a with default parameters [28]. Following alignment, SAMtools (version 1.9) [29] was used to convert the SAM files to binary format (BAM, Binary sequence Alignment Map) sorting of the mapped reads according to chromosome position. Duplicate reads were filtered from the sorted BAM files using the Picard tool’s MarkDuplicates programme (v2.17.11). The single nucleotide polymorphisms (SNPs) were discovered using the Haplotypecaller function of the Genome Analysis Toolkit (GATK, version 3.8). All SNPs were filtered using GATK’s “VariantFiltration” with preliminary filter settings of “QUAL <30.0, QualByDepth (QD) < 2.0, Fisher’s exact test (FS) > 60.0, RMS Mapping Quality (MQ) < 40.0, StrandOddsRatio (SOR) > 3.0, MappingQualityRankSumTest (MQRankSum) < -12.5, ReadPosRankSumTest (ReadPosRankSum) < -8.0>” [30].

In order to gain deeper insights into the distinctive characteristics of Vechur cattle, an in-house script was developed to discern genetic variations based on homozygous conditions in both the dam (MT436) and sire (MT435). Additionally, the cattle Quantitative Trait Loci (QTL) database, specifically release 50, available at https://www.animalgenome.org/cgi-bin/QTLdb/BT/index, was used to investigate the potential correlation between these genetic variations and quantitative traits.

### Copy Number variation(CNV) detection

The read depth-based CNVnator approach [16] was employed to determine genomic CNVs between the vechur sample (MT434) and the ARS-UCD 1.2 bovine reference assembly. According to the author’s recommendations, CNVnator was run on sorted BAM files with a bin size of 100 bp. Following calling, raw CNVs were subjected to quality control to retain confident CNVs. The filtering criteria were P-value < 0.001 (calculated using t-test statistics) and q0 (fraction of mapped reads with zero quality) < 0.5. The genes found in the inferred CNV regions were retrieved and functionally annotated using PANTHER [31].

### Population Structure Analysis

To validate genetic relatedness and population stratification, along with our samples, previously reported data comprising 112 individuals of various Bos indicus breeds were used as reference [18]. These reference populations include Sahiwal (13), Tarparkar (17), Gir (15), Ongole (17), Hariana (18), Kangayam (16), and Vechur (16). Both the data sets were merged using the “vcf-merge” tools of VCFtools [32], and only common SNPs in both data sets were preserved. Then, using the program PLINK (version 1.07) [33], we performed classical multidimensional scaling (MDS) based on pairwise identical-by-state (IBS) distance and rendered the plot using the R package MDS plot.

Linkage pruning was also performed for Admixture analysis using PLINK [33], with parameter--indep-pairwise= 50 10 0.1, which performs linkage pruning with a window size of 50 Kb, window step size of 10 bp, and r2 threshold of 0.1 (i.e. the linkage acceptable threshold). This stage chose a group of independent variants to reduce redundancy. Admixture v1.3.0 [34] was then used to read the PLINK bed file with the default parameters (cross-validation --cv=5) and cluster number (*k*) ranging from two to five. The findings are plotted using R script.

### Screening of Differentially Selected Regions

We employed F_ST_ to detect positive selection signatures in the vechur genome based on whole genome SNPs and other individuals of various Bos indicus breeds were used as reference from previously published data [18]. Firstly, the mean F_ST_ value according to Weir and Cockerham’s pairwise estimator method [35] was calculated using VCFtools (v.0.1.13) [32] in autosomal chromosomes. Genes in the genomic regions with high Z-transformed F_ST_ value (> 7.5) were used to identify their functions in terms of gene ontology. The results of population differentiation were visualized in the form of a Manhattan plot by the qqman R package [36].

## List of abbreviations

BUSCO: Benchmarking Universal Single-Copy Orthologue
BWA: Burrows-Wheeler Alignment programme
FST: Fixation index
GHG: Green House Gases
IBS: Identical-by-state
MDS: Classical multidimensional scaling
NGS: Next Generation Sequencing
OECD-FAO: Organisation for Economic Co-operation and Development and the Food and Agricultural Organization
POA: Partial Order Alignment
QTLs: Quantitative Trait Locus
SNPs: Single Nucleotide Polymorphisms
SNVs: Single Nucleotide variations

## Ethics approval and consent to participate

The samples collected for this study involved prior informed consent of the owners of the cattle, and blood samples were collected by veterinarians using the minimum-harm protocols that apply to blood sampling for routine cattle breeding exercise, disease surveillance and diagnostics.

## Consent for publication

Not applicable.

## Competing interest

The authors declare that they have no competing interests.

## Data availability

The sequence and assembly has been submitted to NCBI with accession: PRJNA957582

## Funding

This project is funded by Kerala Livestock Development Board, Kerala, India and Department of Biotechnology (BT/PR46677)

## Author’s contributions

Concept,Funding-NSS, TVA, TSK, RRP

Animal identification, Sample collection, DNA isolation, sample preparation for sequencing: TVA, RRP, RVM, BB, MK

Data analysis: PVM, NSS, SK Manuscript preparation: NSS, PVM, SK

## Acknowledgements

We thanked Dr. Jose James for his help and encouragement in initiating this project. Dr. N. Sadananda Singh has been supported by Ramalingaswami fellowship(DBT, India) BT/PR46677 (Department of Biotechnology, India), EEQ/2018/001090 (DST, India) and IISER TVM research support. Manpreet was supported by EEQ/2018/001090 (DST india). Dr. Rajesh Vakayil Mani, ^1^, Dr. Balu Bhaskar, Dr. Rajeev Raghavan Pillai and Dr. T. Sajeev Kumar are supported by KLDB. Prof. TV Anilkumar is supported by SCTIMST and IISER TVM. Shivani Kumari and Manpreet Kaur are supported by IISER TVM. Poorvishaa VM is supported by DBT-category I fellowship. Sequencing and initial data analysis was carried out by miBiome, India.

## Notes

### Competing Interest Statement

The authors have declared no competing interest.

### Summary of Updates

1. a spelling error in the the title "varaiation" is replaced with "variation" 2. Figure 1A is updated 3. Figure 2 legend has been updated.

